# NOX4 regulates TGFβ-induced proliferation and self-renewal in glioblastoma stem cells

**DOI:** 10.1101/804013

**Authors:** P García-Gómez, M Dadras, C Bellomo, A Mezheyeuski, K Tzavlaki, A Moren, L Caja

## Abstract

Glioblastoma (GBM) is the most aggressive and common glioma subtype with a median survival of 15 months after diagnosis. Current treatments have limited therapeutic efficacy, thus more effective approaches are needed. The glioblastoma tumoral mass is characterized by a small cellular subpopulation, the Glioblastoma stem cells (GSCs), which has been held accountant for initiation, invasion, proliferation, relapse and resistance to chemo- and radiotherapy. Targeted therapies against GSCs are crucial, and so is the understanding of the molecular mechanisms that govern the GSCs. Transforming growth factor β (TGFβ), platelet growth factor (PDGF) signalling and Reactive Oxygen Species (ROS) production govern and regulate cancer-stem cell biology. In this work, we focus on the role of the NADPH oxidase 4 (NOX4) downstream of TGFβ signalling in the GSCs. NOX4 utilises NADPH to generate ROS; TGFβ induces NOX4 expression, thus increasing ROS production. Interestingly, NOX4 itself regulates GSC self-renewal and modulates Since TGFβ regulates PDGFB in GSC, we analysed how PDGFB modulates NOX4 expression and increases ROS production. Both TGFβ and PDGF signalling regulate GSC proliferation in a NOX4/ROS-dependent manner. The transcription factor NRF2, involved in the transcriptional regulation of antioxidant and metabolic responses, is regulated by both TGFβ and NOX4. This results in an antioxidant response, which positively contributes to GSC self-renewal and proliferation. In conclusion, this work functionally establishes NOX4 as a key mediator of GSC biology.

## INTRODUCTION

Glioblastoma **(GBM)** are the most prevalent and aggressive brain tumors, with a median survival of 15 months. They are characterized by high invasiveness, vascular endothelial proliferation and a strong hypoxic component, which makes GBM refractory to radio- and chemotherapy(Westphal and Lamszus, 2011). A large part of GBM (90%) are primary tumors, while the rest are secondary neoplasms originating through the progression from low-grade gliomas, mainly diffuse or anaplastic astrocytomas(Rinaldi et al., 2016). Current therapeutic strategies include gross total surgical resection followed by treatment with radiotherapy and temozolomide(Stupp et al., 2009). GBM is characterised by intra- and inter-tumoural heterogeneity, high invasiveness and vascular endothelial proliferation. Diversity is apparent at the level of malignant tissue organisation, genomic aberrations and transcript expression, resulting in three different GBM tumour groups – classical, pro-neural and mesenchymal (Verhaak et al., 2010; Patel et al., 2014; Crespo et al., 2015; Wang et al., 2017). Despite major advances in the study and understanding of GBM, the prognosis and treatment are still poor, and this is thought to happen because of the resistance and importance of the cancer-stem-cells (CSCs). The presence of CSCs in GBM, defined as glioblastoma stem cells (GSCs), has been established (Singh et al., 2004); GSCs are characterised by self-renewal capacity, high oncogenic potential, and by being able to generate secondary tumours with the same characteristics as the original one, upon serial xenotranplantation. In vitro, GSCs generate neurospheres, they can differentiate towards neurons, astrocytes and oligodendrocytes, and become less tumorigenic. GSCs are also characterised by the expression of the cell surface protein CD133 (prominin1) and are known to be resistant to several drugs commonly used in therapy regimes (Singh et al., 2003; Bao et al., 2006; Chen et al., 2012). It is widely accepted that GBM originate from the stem cell reservoirs responsible for adult neurogenesis in the brain(Singh et al., 2004; Zhu et al., 2005; Siebzehnrubl et al., 2011; Westphal and Lamszus, 2011). There are several reports that support that neural stem cells localized at the subventricular zone (SVZ) are the origin of glioblastoma (Lee et al., 2018; Gimple et al., 2019). Neural stem cell fate is controlled by environmental cues, among which cytokines play a crucial role (Ming and Song, 2011). Interestingly, the Wnt, Notch, Sonic hedgehog and transforming growth factor β (TGFβ) family signalling pathways control both GSCs and neural stem cells; being all of them involved in brain formation and neurogenesis (Alcantara Llaguno et al., 2011; Ming and Song, 2011).

The TGFβ family is a large group of developmental growth factors, which can be divided into two sub-families. On one hand, the TGFβ sub-family includes TGFβs, a number of the growth and differentiation factors (GDFs), activins and nodal. On the other hand, the bone morphogenetic protein (BMP) sub-family with the anti-müllerian hormone, the BMPs and most of the GDFs. All of these cytokines are multifunctional and their effects will depend on the cellular context, being crucial for organogenesis control, embryonic development and tissue homoeostasis (Moustakas and Heldin, 2009). The TGFβ signalling pathway is mediated by two receptors, the type I and type II. Both receptors form a heterotetrameric complex on the cell surface when the dimeric ligands of the TGFβ family bind (Moustakas and Heldin, 2009). In a sequential mode, the activated ligands interact and bind to the type II receptors, which recruit and phosphorylate the type I receptors. Then, the activated type I receptors recruit and phosphorylate the Smad family proteins at their C-terminal serine residues. In the case of the TGFβ subfamily, the type I receptors phosphorylate Smad2/3, which then assemble into heterotrimeric complexes with Smad4, which translocate the complex into the nucleus to regulate the expression of target genes (Moustakas and Heldin, 2009).

In GBM, elevated TGFβ signalling activity confers poor prognosis to the patient (Rich, 2003; Bruna et al., 2007). GBM bypasses TGFβ-induced cell cycle arrest by the expression of specific transcription factors like forkhead box G1 (FoxG1), which inhibits the TGFβ-mediated expression of p21CIP1 through interaction with FoxO3, thus switching off the suppressive role of TGFβ to promote cell proliferation (Seoane et al., 2004). Several studies demonstrated the involvement of other cytokines interacting with TGFβ in order to promote cell proliferation and inhibit cell cycle arrest and apoptosis; as an example, TGFβ induces the expression of platelet-derived growth factor-B (PDGFB) via Smad2/3 (Bruna et al., 2007; Ikushima et al., 2008), as well as the expression of nodal, another TGFβ family member (Sun et al., 2014), and the activation of the nuclear factor of κ light polypeptide gene enhancer in B-cells (NF-κB) through miR-182 (Song et al., 2012). Apart from inducing cell proliferation, TGFβ is also needed by the GSCs to maintain stemness and self-renewal capacity. TGFβ induces the expression of the cytokine leukemia inhibitory factor (LIF) and the stem cell transcription factor Sox2, via Sox4 and Smad2/3; this increases the self-renewal capacity of the GSCs and their stemness characteristics and enhances the GSC tumour-initiating potential, as proven *in vitro* and *in vivo* (Ikushima et al., 2009; Penuelas et al., 2009).

Oxidative stress and malignancy progression have been correlated in several studies (Egea et al., 2017). ROS are composed mainly by hydrogen peroxide (H_2_O_2_) and oxygen superoxide (O_2_^-^) and their overproduction, which is generated by the reduction of oxygen to hydrogen peroxide, determines oxidative stress. Even though high amounts of ROS can determine cellular damage and/or induce cell death by apoptosis (Simon et al., 2000), recent reports suggest that ROS can act as signalling molecules and can regulate different pathways, especially at low concentrations; this property might depend on the duration to ROS exposure, ROS amount and on ROS location (Sundaresan et al., 1995; Chaudhari et al., 2014). Different signalling pathways have been reported to be activated by ROS, such as ERK1/2, NF-κB, activator protein 1 (AP-1) and TGFβ (Ohba et al., 1994; Lo and Cruz, 1995; Sundaresan et al., 1995; Guyton et al., 1996; Lo et al., 1996; Lakshminarayanan et al., 1998; Junn et al., 2000). Furthermore, some studies claim that ROS have also a role either in stem cell maintenance and differentiation, or in the self-renewal capacity of the neural stem cells (Le Belle et al., 2011; Chaudhari et al., 2014; Kang et al., 2016). ROS production is physiologically highly regulated by the balance between ROS producing and antioxidant agents. One of the main ROS producers are the NADPH oxidases (NOX), which consist of 7 different enzymes. In particular, NOX4 is constitutively active and it does not need any docking site for regulatory subunits (Lambeth and Neish, 2014), NOX4 generates mainly H_2_O_2_ (Takac et al., 2011). In a functional perspective, ROS created by the different NOXs have several functions: they are able to modulate different transcription factors, to inactivate different phosphatases and to help cells migrating via the regulation of the urokinase-type plasminogen activator expression and the regulation of matrix metalloproteases (MMPs) (Holmström and Finkel, 2014; Schröder, 2014). Studies performed in different tumour types, suggested a role of NOX4 in supporting tumour migration and cell survival (Boudreau et al., 2014; Gregg et al., 2014; Li et al., 2015) and they pointed out how NOX4 might play different roles depending on the tumour type. Focusing on GBM, NOX4 and its function has been related to cell growth, survival, invasion and therapeutic resistance by hypoxia-induced radiation (Shono et al., 2008; Hsieh et al., 2012).

In this work, we describe that TGFβ and PDGF induce the expression of NOX4, which causes ROS production. Furthermore, NOX4 has a role in modulating the self-renewal capabilities of the GSCs and their proliferation.

## RESULTS

### TGFβ induces NOX4 in glioblastoma stem cells

In order to study mechanisms by which TGFβ regulates glioma stem cell stemness, we used established GSCs derived from two patients with GBM, U3031MG and U3034MG (Xie et al., 2015). These cells were cultured in stem cell media with or without TGFβ for 24 h, their RNA was extracted and a transcriptomic analysis using the HTA2 Affymetrix platform was performed. TGFβ induced 63 and 52 genes in U3031MG and U3034MG, respectively; while it downregulated 86 and 57 genes in U3031MG and U3034MG, respectively (Figure 1a). In the group of TGFβ-upregulated genes, a subset of 10 genes were common between GSC U3031MG and U3034MG cells (Figure 1b). Among these 10 genes we detected known genes upregulated by TGFβ in GBM, such as LIF and ID1 (Penuelas et al., 2009; Anido et al., 2010). We then focused on NOX4, an enzyme responsible to produce reactive oxygen species, which has been described to contribute to apoptosis, migration, invasion, and differentiation (Egea et al., 2017). The role of NOX4 has been previously studied in the TGFβ signalling pathways in different tumours (Carmona-Cuenca et al., 2008; Jiang et al., 2014), but its role downstream of TGFβ in GBM is still unknown.

**Figure 1.**
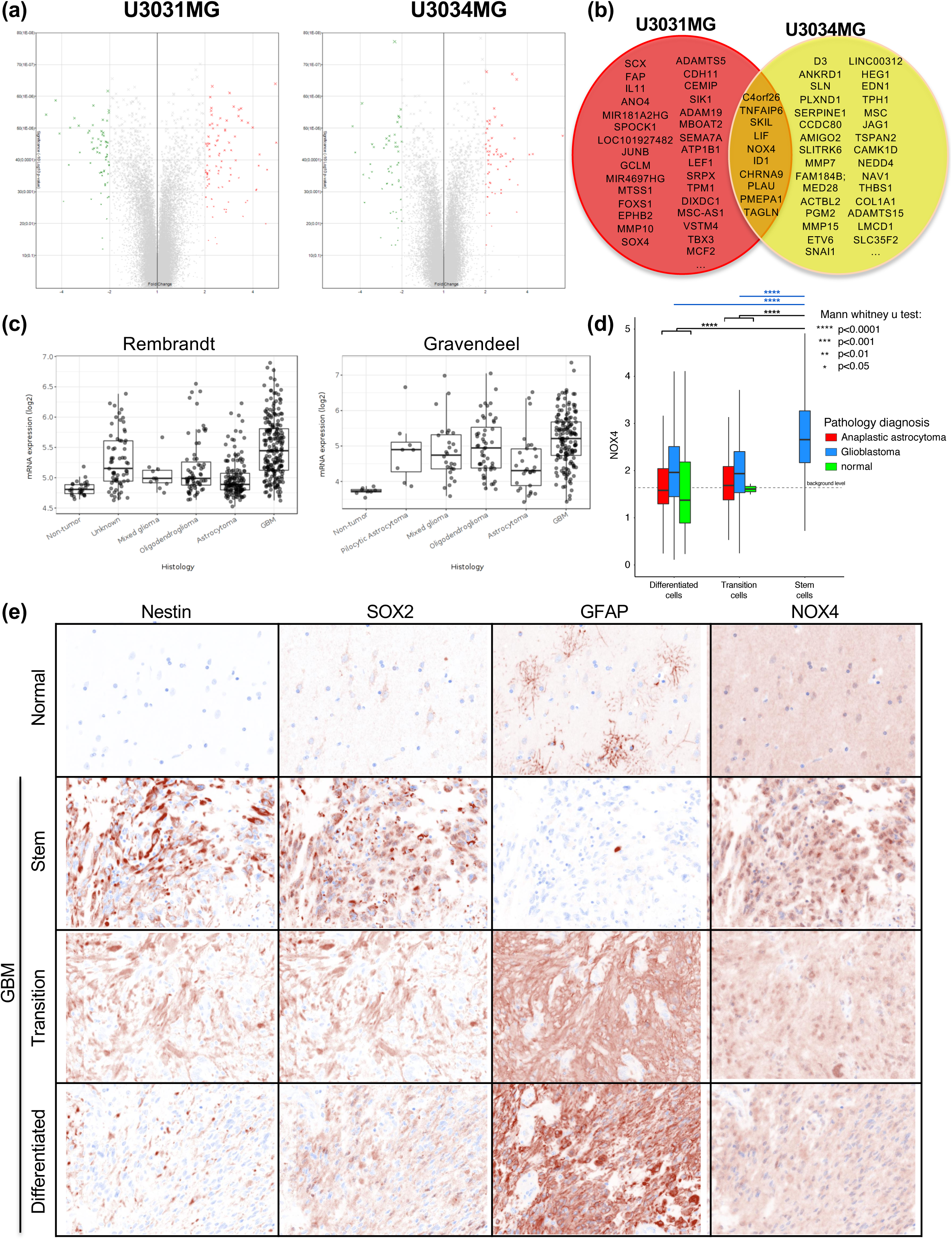
NOX4 is upregulated by TGFβ1 in Glioblastoma Multiforme. (a) Volcano plot of statistical significance against fold-change between untreated and TGFβ1 treated cells in U3031MG cells (left) and U3034MG cells (right). (b) Graphical representation of the common most up-regulated genes in each GSC and their intersection. (c) NOX4 mRNA expression in different types of glioma cells relative to non-tumour cells using different databases (REMBRANDT, left; Gravendeel, right). (d) NOX4 expression per cell in normal, glioblastoma and anaplastic astrocytoma tissue samples (color-coded) plotted as a function of Nestin expression in the same tissue. Cells were classified to three groups, GBM_diff (GFAP^high^/Nestin^low^/SOX2^low^), GBM_transition (GFAP^medium^/Nestin^medium^/SOX2^medium^) and GBM_Stem (GFAP^low^/Nestin^high^/SOX2^high^). Significant differences at \*\*\*\**p*<0.0001. (e) Representative images displaying staining of the four proteins, Par3/Nestin/SOX2/GFAP, in normal brain and glioblastoma tissue samples..

Analysis of NOX4 expression in different glioma subtypes using the Gliovis data portal for visualisation (Bowman et al., 2017) revealed differences between tumour and non-tumour cells, with highest expression of NOX4 recorded in GBM (Figure 1c) in two different datasets, the Repository for Molecular Brain Neoplasia Data (REMBRANDT)(Madhavan et al., 2009) and the Gravendeel dataset (Gravendeel et al., 2009). Correlating this expression with survival expectancy in glioma and GBM, a worse prognosis was recorded in the patients with higher levels of NOX4 compared with patients with low NOX4 expression (Sup. Fig 1a). Moreover, a positive correlation was observed between different TGFβ family members and NOX4 expression (Sup. Fig 1b).

To analyze expression of NOX4 in tissues from GBM patients we performed immunohistochemical analysis on a tissue microarray (TMA) with human GBM, anaplastic astrocytoma and non-tumoral brain samples. In order to analyse whether NOX4 was expresses in GSCs or differentiated tumor cells we performed co-staining with multiple antibodies, aiming at measuring correlations between Par3 protein expression and the GSC subpopulation (Fig. 1d-2). In addition to NOX4, we used antibodies against Nestin and SOX2, which are established stem-cell markers in GBM, and GFAP as an indicator of the astrocytic lineage. We then classified cells in each tumor section into to three groups: GBM_diff (GFAP^high^/Nestin^low^/SOX2^low^) representing more differentiated astrocytes, GBM_transition (GFAP^medium^/Nestin^medium^/SOX2^medium^) possibly representing tumor cells undergoing differentiation transitions, and GBM_Stem (GFAP^low^/Nestin^high^/SOX2^high^) representing the GSCs. Immunostaining of normal brain tissue showed that GFAP was mainly expressed at a moderate level, whereas the stem-cell markers Nestin and SOX2 were almost absent (undetectable, Fig. 1e). Interestingly, in the GBM tissue samples, the GBM cells with high GFAP (GBM-differentiation group) exhibited the lowest NOX4. In contrast, NOX4 expression levels were higher in GBM transition cells and GBM_stem cells (Fig 1d-e).

**Figure 2.**
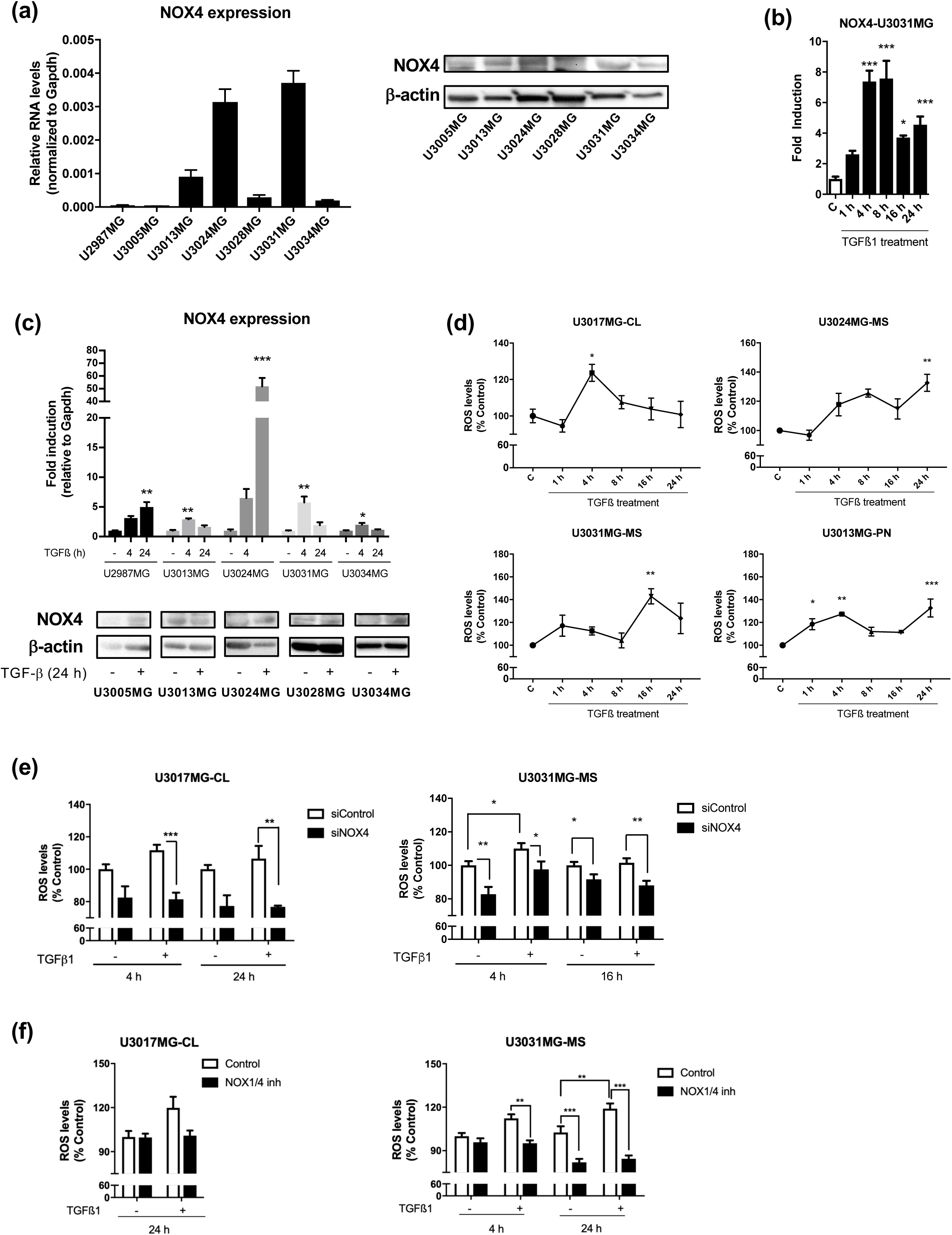
TGFβ1 increases the reactive species of oxygen in a NOX4-dependent manner. (a) Expression panel of *NOX4* in different patient-derived glioblastoma stem cells lines at the RNA level (left) and protein level (right). (b) Time course of NOX4 mRNA expression levels analysed by qPCR in U3031MG cell line upon TGFβ1 treatment, representative experiment. (c) *NOX4* mRNA expression levels analysed by qPCR in different cell lines and time points upon TGFβ1 stimulation. Data represents the mean ± s.e.m. (N = 2-3 biological replicates). (d) Time course of ROS production upon TGFβ1 treatment in different cell lines. Data represents the mean ± s.e.m. (n=2 indenpendent experiments and each with biological triplicate). (e) ROS production upon TGFβ1 stimulation in control (siControl) and NOX4-silenced (siNOX4) cells in different cell lines and time points. Data represents the mean ± s.e.m. (n=3 indenpendent experiments and each with biological triplicate). (f) ROS production upon TGFβ1 stimulation in the presence or absence of NOX1/4 inhibitor in different GSCs cells and time points. Data represents the mean ± s.e.m. (n=2 indenpendent experiments and each with biological triplicate). In the cell line names, CL stands for classical subtype, PN stands for Proneural subtype and MS for mesenchymal subtype of GBM. Statistical comparison indicates * p < 0.05, ** p < 0.01, *** p < 0.001.

### TGFβ stimulates ROS production in a NOX4-dependent manner

NOX4 expression analysis in different GSC cell lines was performed, revealing major differences in NOX4 expression, possibly reflecting individual patient diversity (Figure 2a). TGFβ1 induced the expression of NOX4 at different time points in all GSCs cell lines from distinguished subtypes (Figure 2b-c). Next, we analysed the production of ROS in the GSCs after TGFβ1 treatment, and found that ROS production increased after TGFβ1 at different time points in the different cell lines (Figure 2d). Interestingly, ROS production was reduced either when silencing NOX4 with specific small interfering RNAs (Figure 2e) or when using the enzymatic activity inhibitor of NOX4 (GKT137831) (Figure 2f). It is worth to clarify that GKT137831 inhibits both NOX1 and NOX4, however, NOX1 cannot contribute to ROS production, due to almost undetectable levels of NOX1 in GSCs. In both cases, ROS levels were also diminished to basal or even lower levels when silencing or inhibiting NOX4 in GSCs (Figure 2e-f).

### TGFβ induces self-renewal capacity of GSCs through LIF in a NOX4-dependent manner

In order to study the self-renewal capacity of the GSCs, limiting dilution neurosphere assay was performed. In the neurosphere assay, an increase of the self-renewal capacity of the GSCs was monitored as stem cell frequency: silencing of NOX4 decreased the self-renewal capacity in U3017MG and U3031MG (Figure 3a-b). In accordance to this, silencing of NOX4 diminished the expression of several stem cell markers in U3031MG: CD133/PROM1, OLIG2 and NESTIN (Figure 3c). At the same time, the induction of the astrocytic marker GFAP under differentiation conditions with 10%FBS or BMP7 (60 ng/ml) was enhanced when NOX4 was silenced (Figure 3d). Since previous studies have highlighted the role of autophagy in cancer stem cell maintenance(Vitale et al., 2015), we analyzed the accumulation of autophagosomes by flow cytometry in GSCs after knocking down NOX4 (Supl. Fig. 2a). A decrease in the amount of autophagosomes in U3031MG cells after silencing NOX4 was evident following flow cytometirc analysis, but significant.

**Figure 3.**
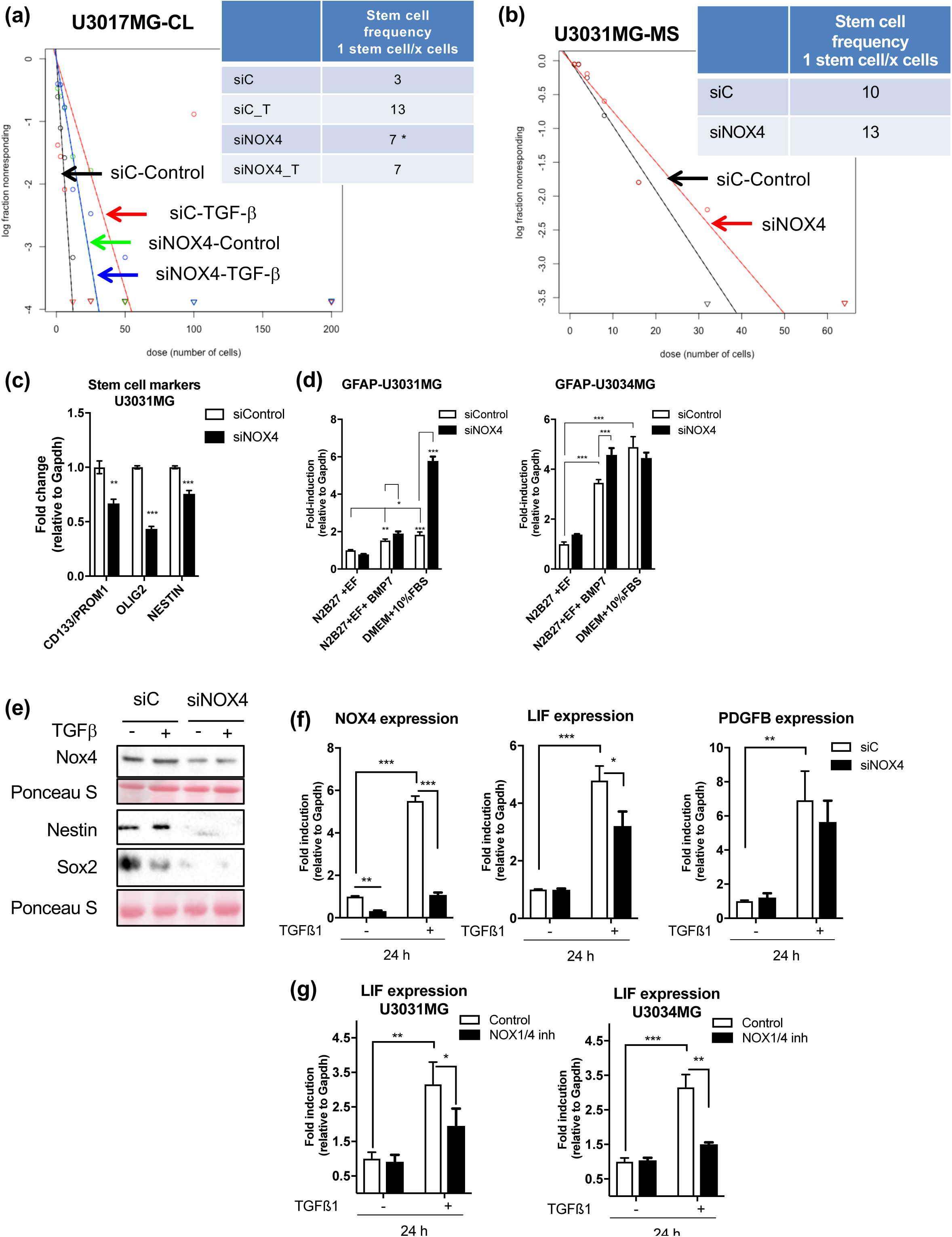
NOX4 regulates GSC self-renewal capacity and TGFβ1-induced stemness. (a, b) U3017MG cells and U3034MG were transiently transfected with control (siControl) or NOX4 (siNOX4) siRNAs and stimulated with TGFβ1, when indicated, for 6 days. Limiting dilution neurosphere assay was performed, analysed by ELDA showing stem cell frequencies. The plot shows the log fraction of wells without spheres as a function of plated cell number. The more vertical the line, the higher the percentage of sphere-forming cells or stem-cells. The tables show the stem cell frequency, p-value < 0.05 was considered as significant. (n = 2 with 12 replicates each). (c) U3031MG cells were transiently transfected with control (siControl) or NOX4 (siNOX4) siRNAs, mRNA expression levels of the indicated genes were analysed by qPCR. Data represents the mean ± s.e.m. (n = 2 independent experiments and each with technical triplicate). (d) U3031MG and U3034MG cells were transiently transfected with control (siControl) or NOX4 (siNOX4) siRNAs, cells were treated with 10% FBS or BMP7 (60 mg/ml) treatment for 3 days to induce their differentiation, mRNA expression levels analysed by qPCR. Data represents the mean ± s.e.m. (n = 2 independent experiments and each with technical triplicate). (e, f) U3031MG cells were transiently transfected with control (siControl) or NOX4 (siNOX4) siRNAs and stimulated with TGFβ1 for 24 h: (e) Immunoblot of the indicated proteins, PonceauS staining is used as a loading control, (f) mRNA expression levels analysed by qPCR. (g) U3031MG or U3034MG cells were stimulated with TGFβ1 for 24 h in with or without NOX1/4 inhibitor, mRNA expression levels analysed by qPCR. Data represents the mean ± s.e.m. (n = 2-3 independent experiments and each with technical triplicate). Statistical comparison indicates * p < 0.05, ** p < 0.01, *** p < 0.001

To understand how TGFβ1 can increase the self-renewal capacity, we analysed the protein expression of Nestin and SOX2, two known stem cell markers: TGFβ1 induced the expression of Nestin but not of SOX2, while the silencing of NOX4 strongly decreased their basal expression (Figure 3e). At the transcriptional level, LIF was upregulated upon TGFβ1 treatment, but this effect was reduced when silencing or inhibiting the function of NOX4 (Figure 3f-g). These results demonstrated a novel role of NOX4 in mediating LIF induction and GSC self-renewal determined by TGFβ signalling. Finally, we also observed that TGFβ1 induced PDGFBB expression, but in a NOX4-independent fashion (Figure 3f).

### PDGF signalling upregulates NOX4 expression and it increases ROS production in a NOX4-dependent manner

Expression analyses were performed using the REMBRANDT and Gravendeel datasets, revealing that the PDGFRB is upregulated in different gliomas, especially in GBM, when compared with non-tumour cells (Figure 4a). Translating this expression pattern to Kaplan-Meier survival expectancy, a worse prognosis was recorded in patients with high expression levels of the PDGFRB (Figure 4b). Finally, the expression of PDGFRB and NOX4 was analysed simultaneously in the patients of the same datasets, observing a significant positive correlation (Figure 4c). Once established the PDGFRB-NOX4 positive correlation and the role of PDGF signalling in the clinical outcome of GBM patients, we studied whether NOX4 expression changed upon PDGFB stimulation. NOX4 was upregulated by PDGFB at different times points in the different cell lines (Figure 4d). We subsequently assessed the ROS production in GSCs upon stimulation with PDGFB, demonstrating that PDGFB was also able to increase ROS production in several cell lines at different time points (Figure 4e), this effect was NOX4-dependent as shown inhibiting NOX4 activity with GKT137831 reduced ROS (Figure 4f).

**Figure 4.**
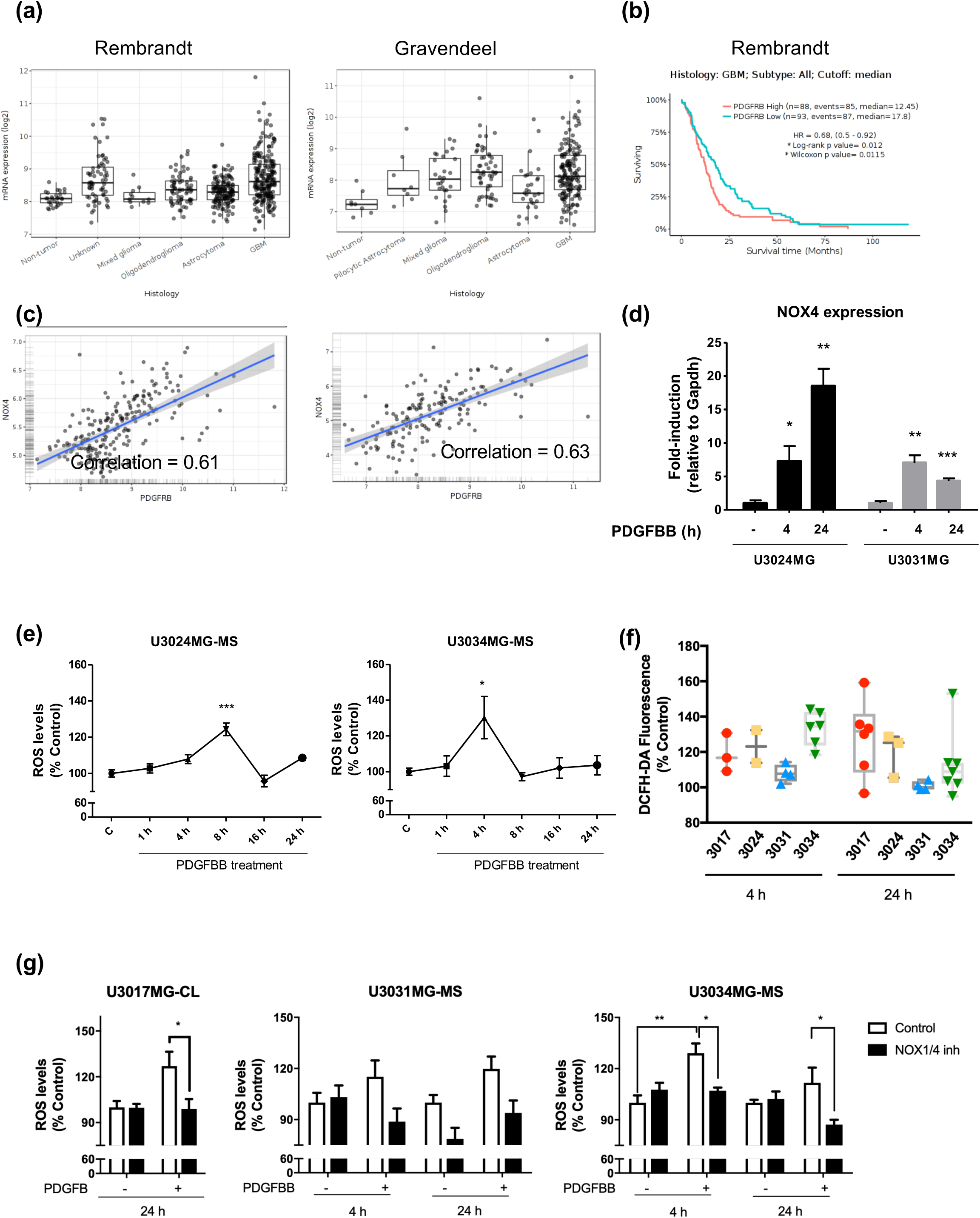
PDGFBB upregulates NOX4 and ROS production in GBM. (a) PDGFRB mRNA expression in different types of glioma cells relative to non-tumour cells using database information (REMBRANDT, left; Gravendeel, right). (b) Kaplan–Meier survival from the Rembrandt database patients according to PDGFRB expression. (c) Scatter plots of NOX4 and PDGFRB expression (Pearson correlation analysis) using normalised mRNA expression data from the REMBRANDT (left) and Gravendeel (right) databases. (d) NOX4 mRNA expression levels analysed by qPCR in different cell lines and time points upon PDGFBB stimulation. Data represents the mean ± s.e.m. (n = 2 independent experiments and each with technical triplicate). (e) ROS production time course upon PDGFBB stimulation in two different cell lines. Data represents the mean ± s.e.m. (n = 1-3 experiments with biological triplicate). (f) ROS production upon PDGFBB stimulation at the indicated time points, data represents the mean ± s.e.m. (n = 1 experiment with biological triplicate). (g) ROS production upon PDGFBB stimulation in NOX4-inhibited cells in different cell lines and time points. Data represents the mean ± s.e.m. (n = 2 experiments with biological triplicate). In the cell line names, CL stands for classical subtype and MS for mesenchymal subtype of GBM. Statistical comparison indicates * p < 0.05, ** p < 0.01, *** p < 0.001

### TGFβ and PDGF increase the proliferation of GSCs in a NOX4-dependent manner

TGFβ and PDGF are known to increase cell proliferation of different cell lineages including glioma cells (Fleming et al., 1992; Seoane et al., 2004; Bruna et al., 2007; Penuelas et al., 2009). Moreover, ROS production determined by NOX proteins has also been positively linked to cell proliferation (Dickinson et al., 2011; Le Belle et al., 2011). In order to understand the role of NOX4 in cell proliferation, a Ki67 immunofluorescence assay was performed in GSCs. The percentage of Ki67-positive nuclei, indicative of actively proliferating cells, was increased when cells were treated with TGFβ1 and PDGFBB while this event was abolished almost to basal levels, when NOX4 was inhibited in both conditions (Figure 5a and 5b). The effect of NOX4 silencing on self-renewal and basal cell proliferation could be due to programmed cell death, as no changes in the percentage of dead cells could be recorded via flow cytometry analysis of the propidium iodide positive cells (Supl. Fig. 2b). These results indicate a crucial function of NOX4 on the proliferative phenotype induced by both TGFβ and PDGF signalling, validating that the NOX4-dependent generation of ROS is physiologically important.

**Figure 5.**
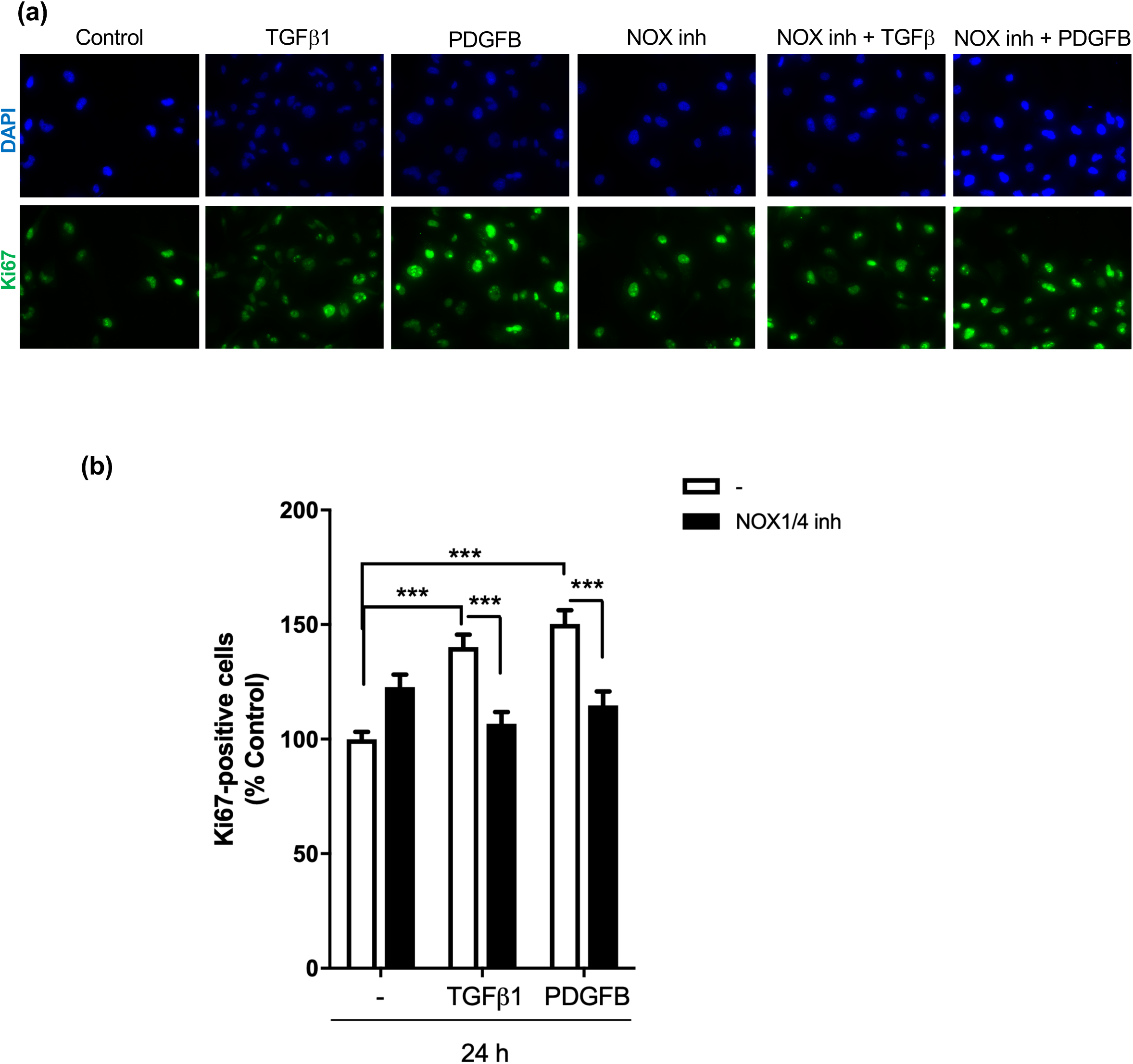
TGF-β1 and PDGFBB increase cell proliferation in a NOX4-dependent manner. U3031MG cells were stimulated with TGFβ1 for 24 h, in the presence or absence of the NOX1/4 inhibitor. (a) Representative microscopic images of immunofluorescence of Ki67 (green) and DAPI (blue). (b) Quantification of Ki67 positive cells with respect to the control upon TGF-β1 and PDGFBB stimulation with or without NOX1/4 inhibitor. Data represents the mean ± s.e.m (n= 3 independent experiment, 10 images per condition and experiments were quantified). Statistical comparison indicates *** p < 0.001

### TGFβ signalling modulates GSC metabolism in a NOX4-dependent manner through the NRF2 pathway

Once elucidated the role of NOX4 in the self-renewal and proliferation of GSCs, we further investigated if the nuclear factor E2-related factor 2 (NRF2) could lie downstream of these NOX4-mediated events, via a potential NOX4-NRF2 axis. NRF2 is a transcription factor known to be regulated by both TGFβ and NOX4 in other cell types, with the capability to reprogram the cellular metabolism in order to support the antioxidant response (Churchman et al., 2009; Brewer et al., 2011). TGFβ1 was able to upregulate the expression of NRF2; however, when NOX4 was silenced or inhibited the induced expression of NRF2 by TGFβ1 was abrogated (Figure 6a), showing a key role of NOX4 in the expression of the NRF2 by TGFβ. Since NRF2 is transcription factor we wanted to analyse whether NOX4 could not only promote its expression but also modulate its transcriptional activity; in order to assess this we used two different luciferase assays: ARE-promoter containing several binding sites of the antioxidant response element (ARE); and the hemoxigenase 1 (HO-1) promoter, a known NRF2 direct target. Both promoters were induced when NRF2 was overexpressed in 293T and HepG2 cells, and their activity was further enhanced when NOX4 was co-expressed with NRF2 (Figure 6b). Furthermore, TGFβ1 induced HO-1 promoter activity, and this was increased when NOX4 was expressed in 293T and HepG2 cells (Figure 6c). NRF2 is known to be a master regulator of cell metabolism, it can contribute to switch from oxidative phosphorylation (OXPHOS) to glycosis (Wang et al., 2018), TGFβ1 induces the expression of one of the glucose transporter, GLUT1, in GBM (Rodríguez-García et al., 2017). We hypothesized that this might be NOX4 dependent. As shown in figure 6d, TGFβ1 was able to increase the expression of Glut1 in a NOX4-dependent manner, as silencing or inhibiting NOX4 reduced GLUT1 expression to basal levels.

**Figure 6.**
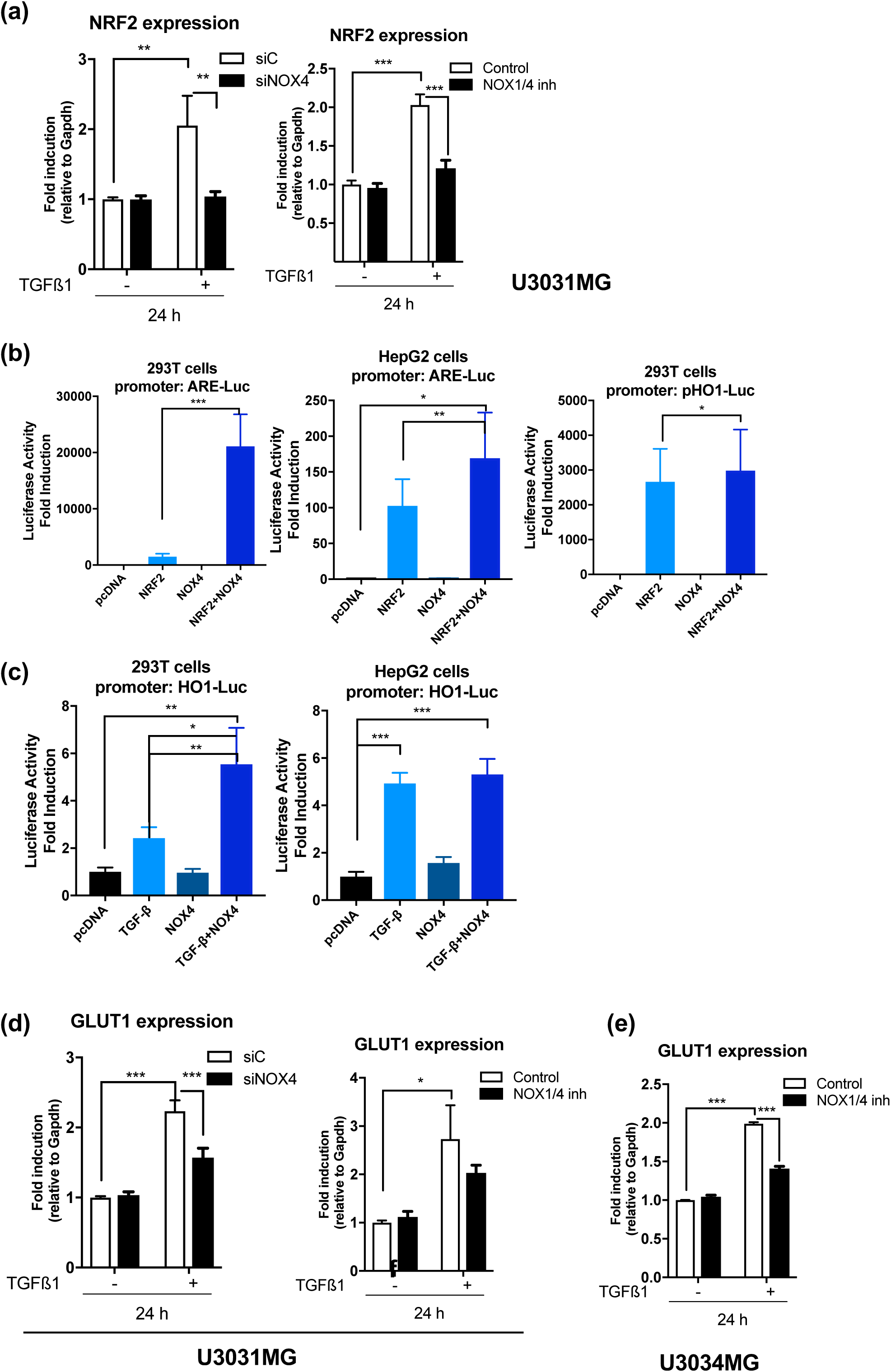
NOX4 has a role TGF-β regulation of NRF2 activity. (a) NRF2 mRNA expression levels analysed by qPCR: U3031MG cells were transiently transfected with control (siControl) or NOX4 (siNOX4) siRNAs and stimulated with TGFβ1 for 24 h (left), or NOX4 function was inhibited by NOX1/4 inhibitor (right). (b) NRF2-responsive ARE-luciferase reporter assay in 293T and HepG2 cells, NRF2-responsive HO1-luciferase reporter assay in 293T cells; cells were transiently transfected with NRF2, NOX4 and the mentioned luciferase reporters, luminescence was measured 48 h after transfection with calcium phosphate. (c) NRF2-responsive HO1-luciferase reporter assay in 293T cells and HepG2 cells; cells were transiently transfected with NOX4 and stimulated for 24 hours prior to the measurement of luminescence. (d) GLUT1 mRNA expression levels analysed by qPCR: U3031MG cells were transiently transfected with control (siControl) or NOX4 (siNOX4) siRNAs and stimulated with TGFβ1 for 24 h (left); U3031MG cells were stimulated with TGFβ1 for 24 h in the presence or absence of the NOX1/4 inhibitor (right). (e) GLUT1 mRNA expression levels analysed by qPCR: U3034MG cells were stimulated with TGFβ1 for 24 h in the presence or absence of the NOX1/4 inhibitor. Data represents the mean ± s.e.m. (n=2 independent experiments and each with technical triplicate). Statistical comparison indicates * p < 0.05, ** p < 0.01, *** p < 0.001

### NOX4 overexpression recapitulated TGFβ1 effects in glioblastoma stem cells

Overexpression of NOX4 in U3034MG, one of the patient-derived GSCs with low endogenous NOX4 expression, resulted in increased intracellular ROS levels (Figure 7a-b); enhanced expression of LIF (Figure 7b) and increased self-renewal capacity (Figure 7d). Interestingly, overexpression of NOX4 resulted in increased expression of the transcription factor NRF2, which might be responsible for the increased expression of GLUT1, GCLM (Figure 7e).

**Figure 7.**
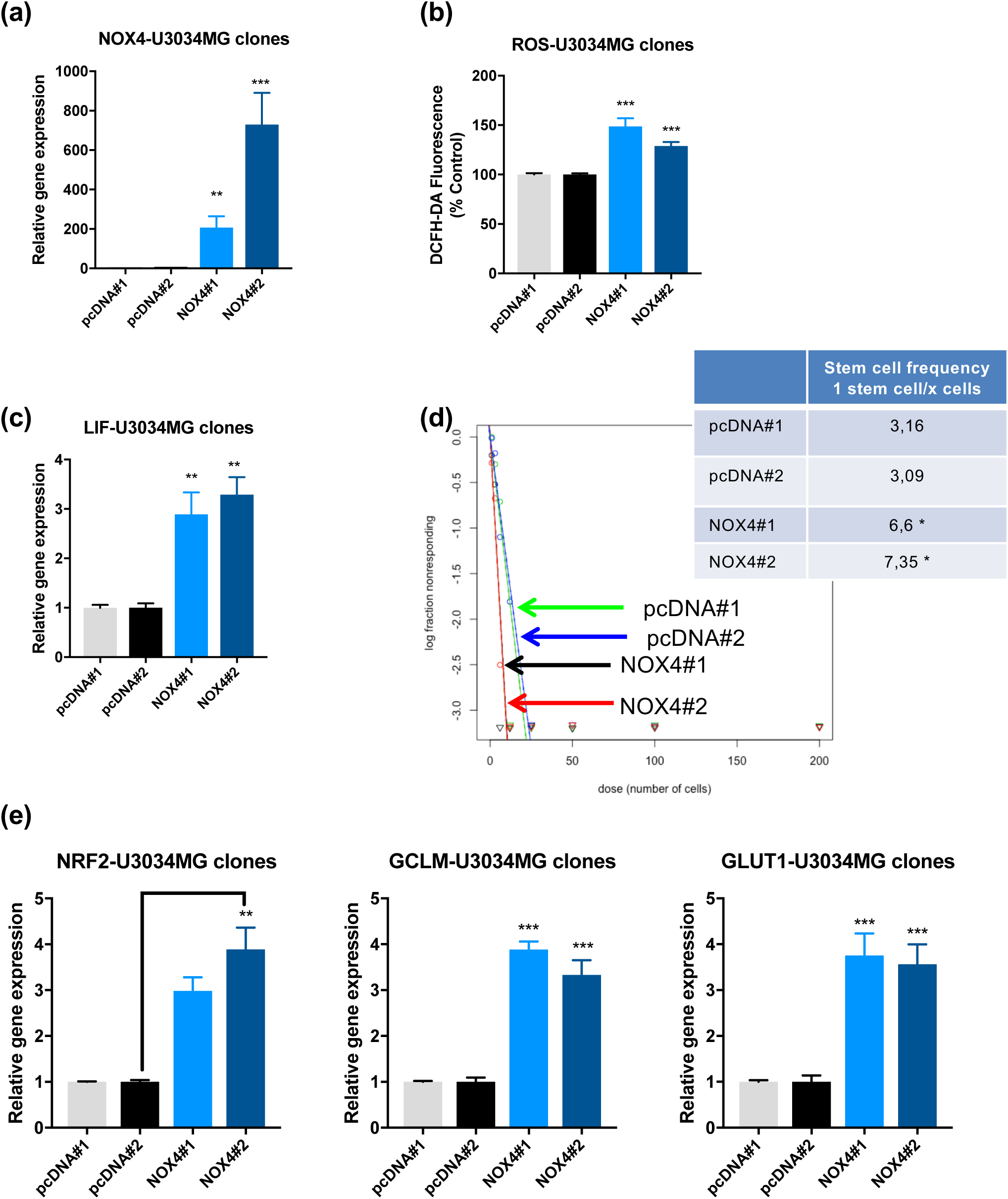
NOX4 overexpression mimics TGFβ effects in GSCs. U3034MG cells stably overexpressing the empty vector pcDNA or pcDNA-V5-NOX4, pool#1 and pool#2 were analysed. (a, c, d) mRNA expression levels analysed by qPCR of the indicated genes, data represents the mean ± s.e.m. (n=2-4 independent experiments and each with technical triplicate). (b) Basal ROS production in the different indicated clones after 48 h of being seeded. Data represents the mean ± s.e.m. (n=3 indenpendent experiments and each with biological triplicate). (d) Limiting dilution neurosphere assay was performed in U3034MG clones in the presence or absence of TGFβ1 for 6 days, analysed by ELDA showing stem cell frequencies. The plot shows the log fraction of wells without spheres as a function of plated cell number. The more vertical the line, the higher the percentage of sphere-forming cells or stem-cells. The tables show the stem cell frequency with lower and upper confidence intervals and the statistical differences between both groups. P. value < 0.05 was considered as significant. (N = 1 with 12 replicates each).

## DISCUSSION

In glioblastoma, one of the most aggressive cancer types, current therapies are not effective and they provide a modest increase in the life expectancy of the patients (Crespo et al., 2015). In this regard, GSCs have emerged as a key element to treat GBM and started to be therapeutic targets (Vescovi et al., 2006) due to their implications in the initiation, migration, invasion, relapse and resistance to chemotherapy of brain malignancies (Bao et al., 2006; Chen et al., 2012). In order to improve the current therapies against gliomas, it is vital to study and to understand the molecular mechanisms that regulate and govern the pathophysiology of GSCs. Several studies identified different signalling pathways involved in glioma, one of them being the TGFβ pathway (Rich, 2003). In GBM, TGFβ is an oncogenic factor, being able to induce proliferation in GSCs, indirectly, by inducing the action of PDGFB, to determine self-renewal, by further upregulating a second cytokine, LIF, and by also causing tumour cell migration, among other functions (Bruna et al., 2007; Penuelas et al., 2009; Anido et al., 2010).

In this work, we observed that TGFβ1 upregulated NOX4 expression in patient-derived GSC cells (Figure 1 and 2), a regulatory event that takes place in other non-tumoral and in cancer cell types such as hepatocellular carcinoma (Carmona-Cuenca et al., 2008; Jiang et al., 2014), with the immediate consequence of increasing the intracellular ROS levels (Figure 2). NOX4 expression levels are increased in GBM compared to lower grade gliomas; and their expression is higher in glioblastoma stem cells compared to differentiated tumor cells (Figure 1). Moreover, high NOX4 expression levels also correlate with worse prognosis of GBM as previously reported (Shono et al., 2008). NOX4 expression correlates with high levels of TGFB ligands, a fact that we address experimentally for the first time.

Neural stem cells are known to have high basal levels of ROS, helping to maintain their self-renewal and proliferation capabilities (Le Belle et al., 2011). In particular, GSCs *per se* had high self-renewal characteristics with high stem cells frequency in the absence of ligand stimulation (Figure 3). NOX4 silencing diminished stem cell frequency and it also reduced the expression of several stem cell markers such as CD133, Nestin and OLIG2. In this paper we confirmed that TGFβ increased the expression of LIF in a NOX4-dependent manner (Figure 3). This suggests a key role of NOX4 and ROS in the TGFβ-dependent self-renewal of GSCs and gives another clue to the possible need for ROS for a proficient TGF-β signal transduction. TGFβ is able to regulate PDGFB (Figure 3) in a Smad2/3-dependent manner (Bruna et al., 2007); but this induction is not NOX4-dependent. Frequently, and especially in GBM, genetic mutations in the PDGFR and over-expression of both receptors and the PDGF ligands, promote tumour generation, proliferation and survival, which emphasises the key role of PDGF signalling in gliomas (Heldin and Westermark, 1999)

PDGF signalling has been reported to depend on ROS in order to generate its signalling cascade (Sundaresan et al., 1995). PDGFB is able to induce both NOX4 expression and to increase intracellular ROS levels, which are produced by NOX4 (Figure 4). Our results suggest a novel regulatory loop where PDGF is able to increase the ROS production. Moreover, TGFβ and PDGFBB stimulate GSCs proliferation, as previously reported by others (Bruna et al., 2007; Lindberg and Holland, 2012); interestingly, their effects on cell proliferation was in fact reduced when inhibiting NOX4 (Figure 5). This opens a new field where both PDGF and ROS-derived signalling can regulate each other and promote different functions in GSCs under the coordinated action of TGFβ.

To understand how NOX4-ROS signalling can regulate self-renewal and proliferation of the GSCs per se and especially downstream of TGFβ signalling, we focused our attention on the transcription factor NRF2. NRF2 is a master transcription factor that is able to modulate cell metabolism in order to support the antioxidant response, enhance the pentose phosphate pathway and fatty acid oxidation, while repressing the lipid metabolic pathway (Vomund et al., 2017). On one hand, NOX4 is known to regulate the transcription factor NRF2 (Brewer et al., 2011); distinctively, TGFβ is also known to transcriptionally regulate this factor (Churchman et al., 2009). We were able to elucidate for the first time that TGFβ-induced NRF2 expression is NOX4 dependent (Figure 6), and that NOX4 promotes NRF2-induced transcriptional activity. TGFβ could be enhancing GSCs proliferation and stemness by increasing glucose uptake by upregulating GLUT1 expression, involving NOX4 (Figure 6)

Finally, overexpression of NOX4 itself is able to recapitulate the effects induced by TGFβ1, such as enhanced self-renewal, and higher levels of LIF and NRF2 which could lead to a metabolic reprogramming of the GSCs, favouring their self-renewal capacity and proliferation. Interestingly, NRF2 expression correlates with stemness in glioblastoma (Zhu et al., 2013).

In conclusion, the data presented in this work demonstrated that both TGFβ1 and PDGFBB signalling pathways are able to regulate and induce NOX4 expression and as a consequence the ROS produced by this membrane oxidase are increased. The NOX4-produced ROS are important for several cellular functions such as proliferation, self-renewal and glucose metabolism. Our data revealed that TGFβ and PDGF pathways regulate these functions via NOX4-derived ROS, opening the field to the possibility of targeting NOX4, as a crossroad node of two major signalling pathways in glioblastoma.

## MATERIALS AND METHODS

### Cell culture

The human GBM cell lines U3017MG-CL, U3024MG-MS, U3031MG-MS and U3034MG-MS (HGCC, Uppsala, Sweden)(Xie et al., 2015), were cultured at 37 °C (5 % CO_2_ and 100 % humidity) under serum-free conditions using the N2B27 medium. In the nomenclature of the cell lines, CL stands for classical GBM subtype, whereas MS stands for mesenchymal GBM subtype. The N2B27 medium was composed by DMEM/F12 (50% v/v), Neurobasal^®^ (50% v/v), B27 without vitamin A (2% v/v), N2 (1% v/v) (Thermo Fisher Scientific, Waltham, MA, US), Glutamine (1% v/v), Penicillin-Streptomycin, (1% v/v) (Sigma-Aldrich Co., St. Louis, Missouri, US); the N2B27 medium was supplemented with EGF (10 ng/mL) and bFGF (10 ng/mL) (Peprotech, Rocky Hill, NJ, US). The culture dishes and plates used (Sarstedt AG & Co, Nümbrecht, Germany) were precoated using poly-ornithine (10 µg/mL in dH_2_O) and laminin (10 µg/mL in PBS) (Sigma-Aldrich Co., St. Louis, Missouri, US).

### Knockdown assays - siRNA transfections

For siRNA-driven knockdowns, cells of 80 % confluency were transfected with 20 nM of ON-TARGETplus SMARTpool human NOX4 siRNA, with a pool of 4 different siRNAs that target several NOX4 transcript variants, and with control siRNA (GE Healthcare Dharmacon, Lafayette, CO, US). The transfections were done using the cationic lipid reagent silentFect^TM^ (Bio-Rad, Hercules, CA, US), according to manufacturer’s instructions. The knockdown efficiency was determined by RT-qPCR at the RNA level.

### Overexpression Assays – Stable Clone generation

Different stable clones were created using U3034MG-MS cell line with pcDNA empty plasmid and pcDNA-V5-NOX4, which was kindly supplied by Professor Ulla Knaus (University College of Dublin, Ireland). The plasmids were transfected into U3034MG-MS cells. To do so, 6-well plates with 80% confluent cells were transfected by using TransIT-X2^®^ Dynamic Delivery System (Mirus, Madison, WI, US), according to manufacturer’s instructions. Concisely, 2.5 µg of plasmid DNA were mixed and incubated with TransIT-X2 in OptiMEM^®^ I Reduced Serum Medium (Thermo Fisher Scientific, Waltham, MA, US) for 30 min at RT to create DNA-TransIT-X2 complexes, in a ratio of 3 µL TransIT-X2/µg of DNA. Finally, the complexes were added to the cells with N2B27 medium (2.5 mL) by drops and incubated at 37 °C (5 % CO2 and 100 % humidity) for 48 h. After incubation, the medium was changed adding 1 mg/ml of Geneticin (G418) (Thermo Fisher Scientific, Waltham, MA, US), which is a selective agent for eukaryotic cells in order to select the positive transfected cells (10 mg/mL in N2B27 medium).

### Intracellular Reactive Oxygen Species (ROS) measurements

Intracellular ROS measurements were performed in order to evaluate its levels after stimulation or inhibition with the different growth factors or inhibitors, by using a fluorimetric method. Cells were stimulated with TGFβ1 (5 ng/mL) (Peprotech, Rocky Hill, NJ, US), PDGFBB (20 ng/mL) (Ray Biotech, Norcross, GA, US); and/or NOX1/4 inhibitor (GKT137831) (20 µM) (Cayman Chemical, Ann Arbor, Michigan, US), the inhibitor was added 30 minutes prior stimulation. In a previously coated 12-well plate, 100,000 cells were seeded per well and allowed to attach by incubating them 24 h at 37 °C (5 % CO_2_ and 100 % humidity). In each experiment, every condition was performed in 3-4 technical replicates. At the collection time, the cells were washed with PBS and incubated in dark with 2’,7’-dichlorodihydrofluorescein diacetate (H_2_DCF-DA) (2.5 µM in HBSS) (Thermo Fisher Scientific, Waltham, MA, US) for 30 min at 37 °C (5 % CO_2_ and 100 % humidity). After incubation, the cells were lysed with 250 µL of a specific lysis buffer composed by HEPES (25 mM and pH 7.5), MgCl_2_ (1.5 mM), NaCl (60 mM), EDTA (0.2 mM) and Triton X-100 (1 % v/v) (Sigma-Aldrich Co., St. Louis, Missouri, US) while shaking at 4 °C during 20 min. In duplicate (100 µL), the lysed cells were transferred to a 96-well plate. Fluorescence was measured using an excitation and emission wavelength of 485 and 520 nm, respectively in the EnSpire^®^ Multimode Reader (Perkin Elmer, Waltham, MA, US). The 50 µL left were used for protein quantification using BCA (Material and Methods 4.4). The results were expressed as a percentage of control (% Control) and calculated as fluorescence units per µg of protein.

### RNA extraction, quantitative reverse transcription polymerase-chain-reaction (RT-qPCR) and HTA2 affymetrix platform array

In previously coated 6-well plates, 200,000 cells were seeded and allowed to attach by incubating them 24 h at 37 °C (5 % CO_2_ and 100 % humidity). Cells were treated with DMSO (1% v/v), TGFβ1 (5 ng/mL), PDGFBB (20 ng/mL), NOX1/4 inhibitor (GKT137831) (20 µM). Total cellular RNA was extracted from the cells by using NucleoSpin^®^ RNA Plus kit (Macherey-Nagel GmbH & Co. KG, Düren, Germany) according to manufacturer’s instructions. Equal amounts of total RNA were reverse-transcribed using iScript™ cDNA Synthesis Kit (Bio-Rad, Hercules, CA, US) according to manufacturer’s instructions. Quantitative PCR was performed using the created cDNA in triplicates by using the CFX Connect™ Real-Time System and CFX Manager (Bio-Rad, Hercules, CA, US) and KAPA SYBR^®^ FAST qPCR Kit (Kapa Biosystems, Wilmington, MA, US) according to manufacturer’s instructions. *GAPDH* was used as a reference gene. The following DNA primers (Table 1) were used.

**Table 1.**
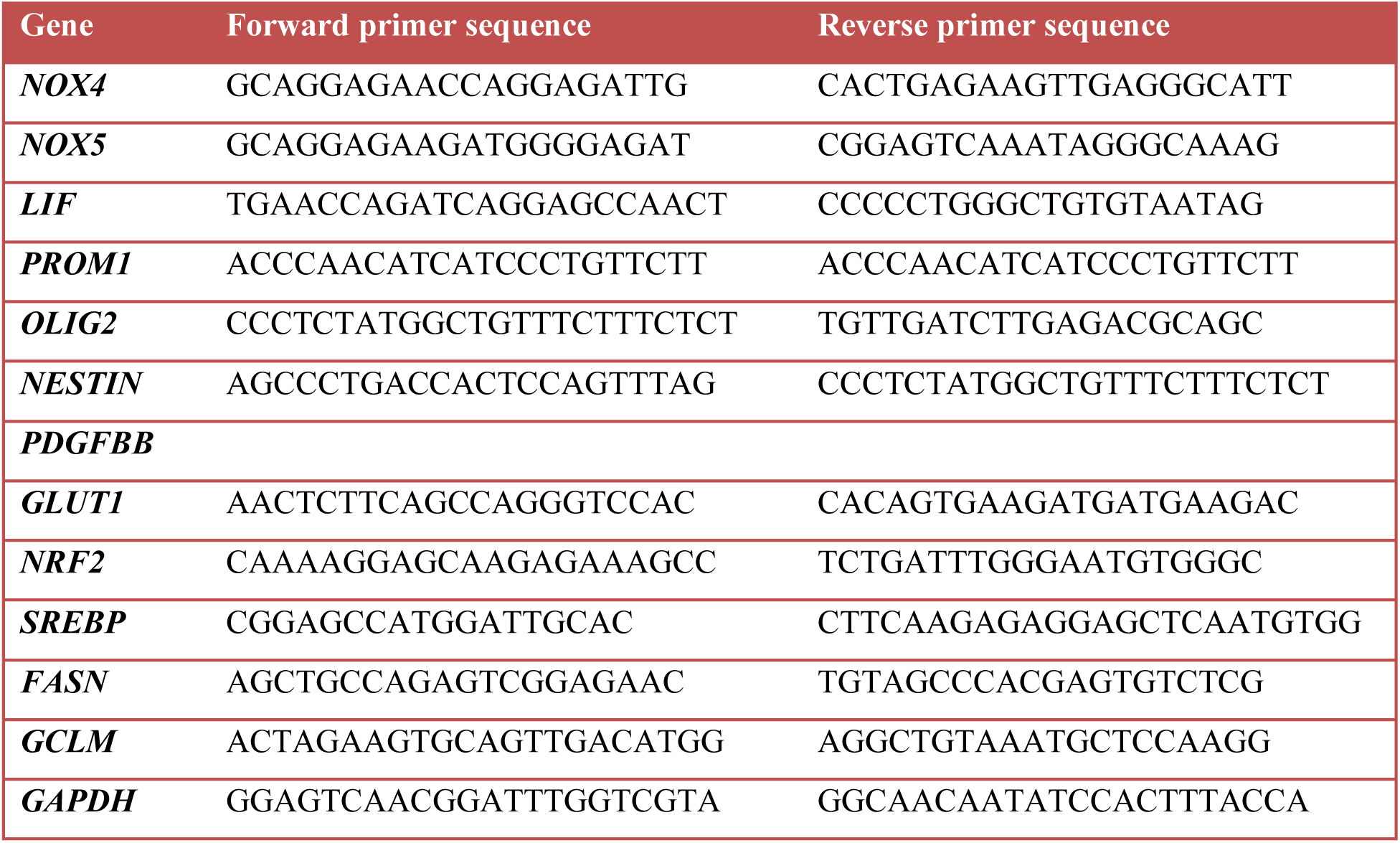
Forward and reverse primer sequences (5’ to 3’) used for qPCR.

Transcriptomic analysis was performed by an HTA2 Affymetrix Platform array (Thermo Fisher Scientific, Waltham, MA, US). For each condition, triplicates were analysed by the Swegene centre for Inegrative Biology at Lund University (SCIBLU). Basic Affymetrix chip and Experimental Quality Analyses were performed using the Expression Console Software v1.4.1.46; differential gene expression analysis was done with Transcriptome Analysis Console (TAC). Adjusted *p*-values (padj) for multiple testing, using Benjamini-Hochberg to estimate the false discovery rate (FDR), were calculated for final estimation of differential expression (DE) significance. The expression profiles have been deposited to Array Express and accession number E-MEXP-xxxx will soon be obtained.

### Luciferase assay

HepG2 or 293T cells (1.8 × 10^4^ cells per 24-well) were transfected with luciferase-encoding together with pCMV-β-galactosidase plasmids (100 ng), the latter as reference, using Lipofectamine-3000 (Life Technologies, Stockholm, Sweden) for 48 h, and assayed as described. The plasmids were: synthetic NRF2-binding promoter ARE-luc, *hHO1* promoter-luciferase, pEF-NRF2 (these plasmids were a kind gift from Prof. Ken Itoh (Hiroshaki University, Japn), and the plasmid pCDNA3-V5-NOX4 from Prof. Ulla Knaus (University College of Dublin, Ireland). Cells were stimulated with TGFβ1 (5 ng/ml) for 24 h. 2 independent biological experiments were performed in 3 technical replicates per condition.

### Immunoblot

Western-Blot assays were performed to observe differences at the protein level among the different conditions. At the collection time, the media was removed and cells were scraped with PBS and collected into a tube by centrifugation at 600 g during 5 min. Then, the pellet was resuspended in 200 µL of lysis buffer and sat on ice for 20 min. Afterwards, a centrifugation at 4 °C at 15800 g during 15 min was done and the supernatant was collected. The lysis buffer was composed by 0.5 % v/v Triton X-100, 0.5 % m/v sodium deoxycolate, 10 mM EDTA, 20 mM tris(hydroxymethyl)aminomethane (Tris) at pH 7.4 and 150 mM NaCl (Sigma-Aldrich Co., St. Louis, Missouri, US) in dH_2_O. Proteases inhibitors and phosphatase inhibitors were also added, cOmplete™, EDTA-free Protease Inhibitor Cocktail (Roche, Basel, Switzerland) and PhosSTOP™ (Roche, Basel, Switzerland). The protein concentration was measured by BCA.

Proteins were analysed by sodium dodecyl sulphate (SDS) polyacrylamide gel electrophoresis (PAGE) and detected by immunoblotting. The primary antibodies against the following proteins were used: NOX4, kindly supplied by Isabel Fabregat (Caja et al., 2009) (1:1,000, rabbit); NESTIN (1:2000 T, rabbit) (ab105389, Abcam^®^, Cambridge, UK), and SOX2 (AB5603, Millipore, Solna, Sweden). For loading controls, the following antibodies were used GAPDH (1:10,000 in TBS-T, mouse) (sc69879, Santa Cruz Biotechnology, Heidelberg, Germany)

### Extreme Limiting Dilution Assay

For measuring the self-renewal capacity of the cells, a limiting dilution neurosphere assay was performed. Briefly, cells were seeded in Corning^®^ Costar^®^ Ultra-Low attachment 96-well plate (Corning Incorporated, Corning, NY, US) performing a serial dilution 1:2 from 200 to 1 cell, performing 12 replicates in each case. The cells were treated with TGFβ1 (5 ng/mL) and control solution (0.1 % BSA) in N2B27 medium and incubated for 7 days at 37 °C (5 % CO_2_ and 100 % humidity). After the incubation time, the wells with neurospheres > 50 µm were counted as positive. The neurospheres were visualised using a phase contrast Axiovert 40 CFL microscope (Carl-Zeiss, Oberkochen, Germany). The data was processed by the tool Extreme Limiting Dilution Analysis (ELDA) available online (Hu and Smyth, 2009).

### Immunofluorescence microscopy

An immunofluorescence assay was performed to analyse cell proliferation. In previously coated 8-chamber polystyrene-treated culture chamber slides (Falcon^®^, Corning, NY, US) 20,000 cells were seeded and allowed to attach during 24 h at 37 °C (5 % CO_2_ and 100 % humidity). Afterwards, the cells were treated with DMSO (1% v/v), TGFβ1 (5 ng/mL), PDGFBB (20 ng/mL) and/or NOX1/4 inhibitor (GKT137831) (20 µM) during 24 h in N2B27 medium. After the treatments, the cells were washed with PBS and fixed with 3.7 % (w/v) paraformaldehyde (PFA) in PBS for 15 min. Subsequently, fixed cells were washed three times with PBS and blocked with 10 % fetal bovine serum (FBS) in PBS with 1 % BSA for 1 h at RT to avoid nonspecific binding. Washing step was performed with PBS. The fixed cells were permeabilised with 0.1% Triton-X-100 in 0.1 % BSA. Then, the cells were incubated at RT for 2 h with primary antibodies against Ki67 (1:1,000 in PBS containing 1 % BSA, rabbit) (ab15580, Abcam^®^, Cambridge, UK). Afterwards, 3 washes steps with PBS were performed and the cells were incubated for 1 h in dark at RT with donkey anti-Rabbit IgG Alexa Fluor 488 secondary antibody (1:200 in PBS with 1 % BSA) (A21206, Thermo Fisher Scientific, Waltham, MA, US). Then, cells were incubated in dark at RT for 5 min with DAPI (1:1,000 in PBS with 1 % BSA) (Sigma-Aldrich Co., St. Louis, Missouri, US). Later, cells were washed 3 times with PBS and the cover slips were mounted on the slides using Fluoromont-G^®^ mounting medium (Southern Biotech, Birmingham, AL, US) and dried in dark at RT. The cells were acquired under the fluorescence microscope Eclipse 90i and processed using NIS-Elements software (Nikon, Tokyo, Japan).

The quantification of the proliferative phenotype of the cells after treatments was performed by quantifying the Ki67-positive cells, by counting the Ki67-positive signal inside the nuclei of the cells by using Fiji image software (Schindelin et al., 2012). The results were expressed as a percentage of proliferation against the control (% Control).

### Cell death measurement

Cell death analysis was performed by quantifying propidium iodide (PI) incorporation by fluorescence-activated cell sorter (FACS) analysis. U3031MG cells were transiently transfected with control (siControl) or NOX4 (siNOX4) siRNAs and stimulated with TGFβ1 for 24, adherent cells and floating dead cells were harvested using accutase and centrifuged for 5 min. The cells were then resuspended in a final volume of 300 µl of PBS plus 0.5 µg/ml of PI. Apoptosis was analyzed in a BD Accuri CG Plus flow cytometer (BD Biosciences, Stockholm, Sweden).

### Autophagy measurement with CytoID

To measure the number of autophagic vacuoles, the Cyto-ID autophagy detection kit (Enzo Life Sciences, Solna, Sweden) was used. Cells were transfected with control and Par3 siRNAs, and the following day cells were treated with chloroquine for the last 16 h of the experiments. Two days after transfection, cells were harvested with accutase, washed and stained for 30 min with Cyto-ID dye, and analysed using a BD Accuri CG Plus flow cytometer (BD Biosciences, Stockholm, Sweden). Quantification was done by using FlowJo software version 10.4.2.

### Multiplex immunohistochemical staining

We analyzed a TMA that contained 35 samples of independent glioblastoma patients and 5 samples of normal brain tissue, each in duplicate, generating 80 tissue cores per TMA (TMA GL806d; US Biomax, Derwood, MF, USA). Mutiplexed immunohistochemical analysis was previously described with the following adaptations (Mezheyeuski et al., 2018). TMA slides were deparaffinized in xylene, hydrated in graded alcohols, and fixed in neutral buffered formalin for 20 min. For antigen retrieval, the slides were boiled in pH 6.0 buffer (AR6001, PerkinElmer Sverige AB, Upplands Väsby, Sweden) for 15 min at 100 °C, using a microwave oven. The primary antibodies used were: rabbit anti-NOX4, kindly supplied by Isabel Fabregat (1:100), rabbit anti-Nestin (1:100; Millipore/Merck, Stockholm, Sweden), rabbit anti-GFAP (1:50; Santa Cruz Biotechnology Inc., Santa Cruz, CA, USA) and rabbit anti-SOX2 (1:100; Abcam, Cambridge, UK). After the first staining, the slides were boiled again to remove primary and secondary antibodies, followed by staining for the second antigen. The primary antibodies were diluted (1:100 or 1:50) in ready-to-use antibody diluent and blocking buffer (ARD1001EA, PerkinElmer Sverige AB, Upplands Väsby, Sweden) and slides were incubated for 30 min at room temperature, followed by incubation with anti-rabbit/mouse Opal Polymer HRP ready-to-use immunohistochemistry detection reagent (ARH1001EA, PerkinElmer Sverige AB, Upplands Väsby, Sweden) for 10 min and Opal 570 and Opal 650 (FP1488001KT and FP1496001KT, PerkinElmer Sverige AB, Upplands Väsby, Sweden) fluorophores for 10 min at room temperature. Afterwards, DAPI staining and mounting with ProlongTM Diamond Antifade Mountant (Thermo Fisher Scientific, Rockford, IL, USA) completed the protocol.

Using the Vectra Polaris system (PerkinElmer Sverige AB, Upplands Väsby, Sweden) multispectral imaging mode, the TMA cores were scanned at 20× magnification. Image analysis was performed by the Inform software (PerkinElmer multispectral imaging mode) by applying spectral unmixing, cell segmentation and recording the mean expression level of each stained antigen in every cell region. Visual evaluation was used to define intensity thresholds that separated specific signals from background. Imaging data analysis was performed by the R software, version 3.3.3(Team R, 2015; RC, 2017). All cells were split into three classes of tumor tissue according to Nestin expression: Nestin signal below background threshold (differentiated cells), Nestin signal above background threshold and below the mean intensity level (transition cells) and Nestin signal above the mean intensity level (stem cells). Par3 intensity was then normalized by setting the background threshold to ‘zero’ intensity. Cells with Par3 signal below background threshold were excluded from analysis, resulting in 55,952 cells used for analysis. Statistical difference between Par3 levels in different cell subgroups was calculated by the Wilcoxon signed rank test for two samples (antibodies), which is equivalent to the Mann-Whitney U-rank test.

### Bioinformatic analysis

Different bioinformatics analyses about gene expression, survival analysis and different genes correlation were performed using different clinical databases with GBM specific data: Gravendeel (n = 159 and n = 284 when using all cancer types of glioma) (Gravendeel et al., 2009) and Rembrandt (n = 203) (Madhavan et al., 2009). The data visualisation tool for Brain Tumour datasets Gliovis was used (Bowman et al., 2017). In the expression correlation studies, the Pearson’s correlation was used. The data was obtained from the datasets on 25/10/2017.

### Statistics

We thank Aristidis Moustakas for reagents and advice, Aslam Aref and Natsnet Tekleab for technical assistance, and laboratory members for useful discussions. The statistical analysis was performed by using the software GraphPad Prism^®^ v.7 (GraphPad Software, San Diego, California, US). The data is presented as the average ± standard error of the mean (SEM). The statistical tests used were one-way or two-way ANOVA followed by Dunnett’s post-test or Bonferroni post-test. A p-value < 0,05 was considered as statistically significant. The significance was considered as * p < 0.05, ** p < 0.01 and ***p < 0,001, as it is indicated in each figure.

## ACKNOWLEDGMENTS

HTA2 microarray analysis analysis was performed at the Swegen Centre for Integrative Biology at Lund University, Sweden. This work was supported by the Ludwig Cancer Research, and by postdoctoral fellowships (LC) from Lisa Erikssons minnesfond (2013) and the Swedish Cancer Society (CAN 2012/1186). OE och EDLA Johanssons stifelse, Petrus och Augusta Hedlunds Stiftelse (M2019-1065), and Svenska Läkaresällskapets fonder (SLS-887701)

## AUTHOR CONTRIBUTIONS

L.Caja conceived the project. L. Caja and P. García-Gómez designed the experiments. L. Caja, P. García-Gómez, M. Dadras, C. Bellomom, K. Tzavlaki, A.Moren acquired the data. L. Caja, P. García-Gómez, and A. Mezheyeuski analyzed the data. L. Caja and P. García-Gómez interpreted the data. L. Caja and P. García-Gómez drafted the article. All authors critically revised the article for important intellectual content and provided final approval prior to submission for publication.

---

**Supplementary Figure 1.**
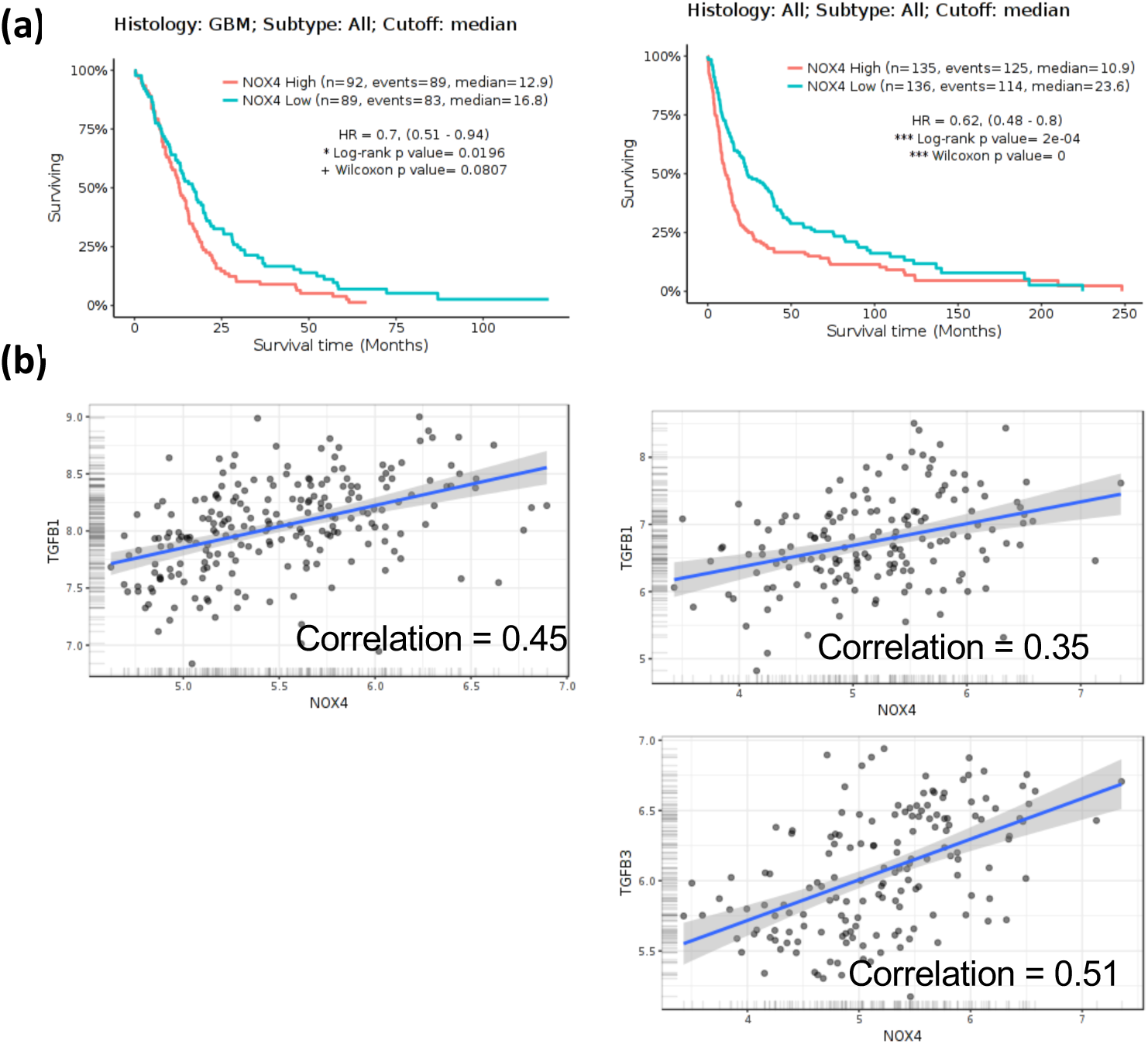
High NOX4 expression correlates with worse prognosis. (a) Kaplan–Meier survival according to NOX4 expression, (REMBRANDT, left; Gravendeel, right). (b) Scatter plots of *NOX4* relative to *TGFB1* and *TGFB3* expression (Pearson correlation analysis) using normalised mRNA expression data from REMBRANDT database (left) and Gravendeel database (right).

**Supplementary Figure 2.**
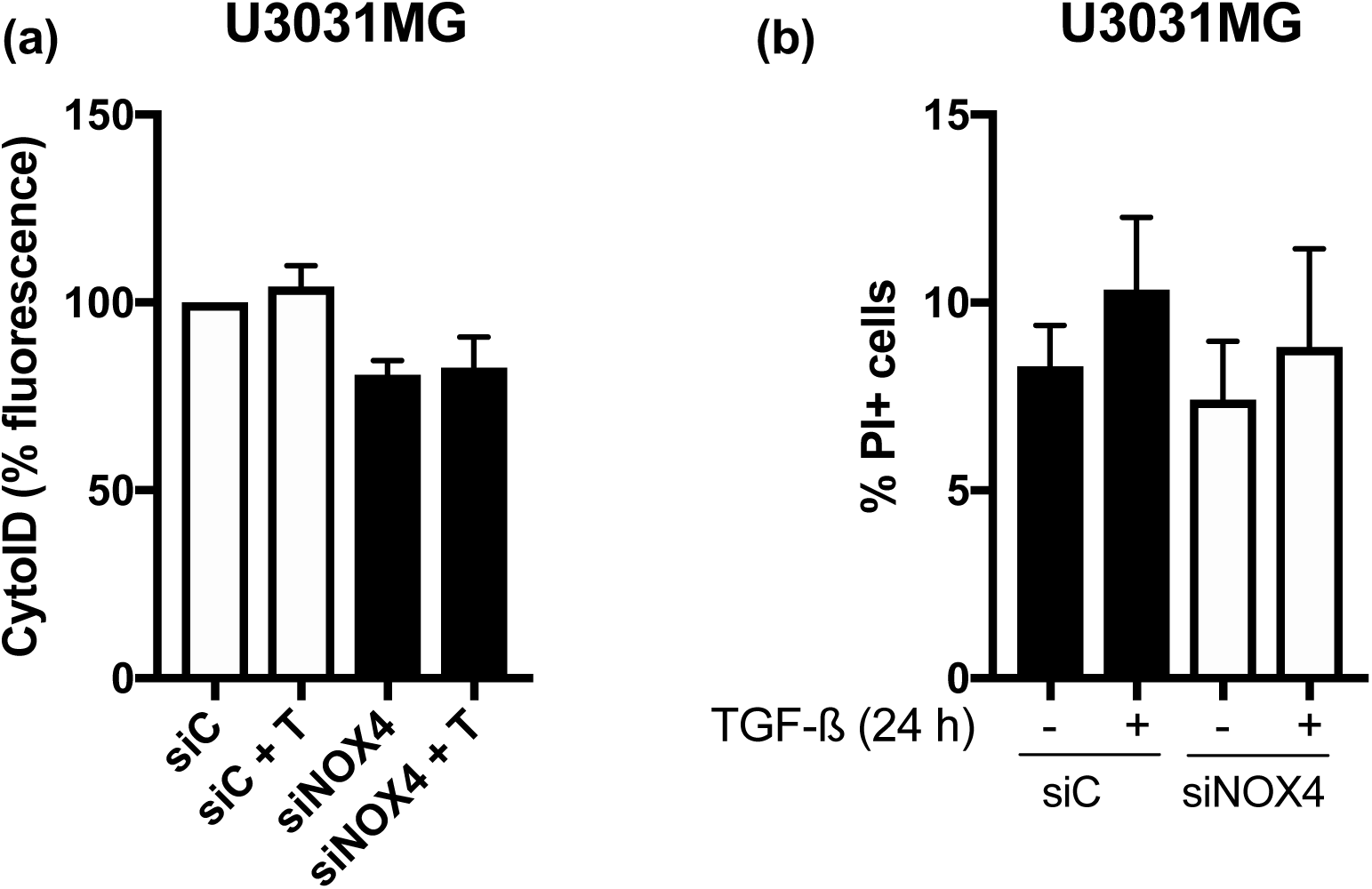
NOX4 silencing does not induce apoptosis. U3031MG cells were transiently transfected with control (siControl) or NOX4 (siNOX4) siRNAs and stimulated with TGFβ1 for 24 h. (a) Measurements of autophagosomes by flow cytometry using cytoID results are mean±S.E.M. of 4 independent experiments. (b) Apoptosis was measured by PI staining, mean±S.E.M. of 4 independent experiments.

